# hRUV: Hierarchical approach to removal of unwanted variation for large-scale metabolomics data

**DOI:** 10.1101/2020.12.21.423723

**Authors:** Taiyun Kim, Owen Tang, Stephen T Vernon, Katharine A Kott, Yen Chin Koay, John Park, David James, Terence P Speed, Pengyi Yang, John F. O’Sullivan, Gemma A Figtree, Jean Yee Hwa Yang

## Abstract

Liquid chromatography-mass spectrometry based metabolomics studies are increasingly applied to large population cohorts, running for several weeks to months, even extending to years of data acquisition. This inevitably introduces unwanted intra- and inter-batch variations over time that can overshadow true biological signals and thus hinder potential biological discoveries. To date, normalization approaches have struggled to mitigate the variability introduced by technical factors whilst preserving biological variance, especially for protracted acquisitions. Here, we designed an experiment with an arrangement to embed biological sample replicates to measure the variance within and between batches for over 1,000 human plasma samples run over 44 days. We integrate these replicates in a novel workflow to remove unwanted variation in a hierarchical structure (hRUV) by progressively merging the adjustments in neighbouring batches. We demonstrate significant improvement of hRUV over existing methods in maintaining biological signals whilst removing unwanted variation for large scale metabolomics studies.

## Introduction

Liquid chromatography-tandem mass spectrometry (LC-MS/MS) is a preferred method of metabolomic acquisition given its high sensitivity and dynamic range. Typically, a range of metabolites can be separated on a single high performance LC column and their relative abundance quantified in MS/MS. This enables capture of fingerprints of specific biological processes that are critical in precision medicine applications such as studying complex metabolic diseases, and discovering new therapeutic targets and biomarkers^1^. There are a number of large-scale cohort studies that have performed metabolomic analyses, such as the Consortium of Metabolomics Studies (COMETS)^2^, and the Framingham Heart Study (FHS)^3^.

Despite a rapid increase in the number of large-scale metabolomics studies, the normalization of metabolomics data remains a key challenge^4^. Due to the data acquisition time of studies with large sample size, prolonged study recruitment and potentially multiple samples at various time points for each participant, the data acquisition process may require the samples be divided into multiple batches, and may span anywhere from months to years^4,5^. Signals often drift over extended periods due to multiple factors including buffer changes, pooled quality control (QC) sample solutions, instrument cleanliness, and machine scheduled maintenance^6^. Common intra-batch variations include changes in LC-MS/MS performance due to instrument-dependent factors such as component failure or inconsistency, and fouling of the column, LC or MS source. Common inter-batch variations include time-dependent instrument variations such as instrument cleaning, tuning, column change, or inconsistent sample preparation factors including change in equipment and operator. These technical factors have substantial impact in downstream analytics and need to be appropriately accounted for to maximise the opportunity to identify true biological signals.

Several workflows have been designed for analysing metabolomics data (e.g. MetaboAnalyst^7^ and NormalyzerDE^8^). However, most of them adapt common normalization methods developed for other omics platforms and do not account for signal drift across extended time or inter-plate variations which are distinct unwanted variations commonly observed in metabolomics studies. A limited number of metabolomics specific normalizations methods have been developed (Table 1). These include: Support Vector Regression (SVR)^5^, Systematic Error Removal using Random Forest (SERRF)^9^, and Removal of Unwanted Variation based approaches^10,11^. These approaches utilise pooled QC samples to control signal drift by fitting non-linear models or by estimating inter-plate variations. The common assumption of these approaches is that the pooled QC samples are identical over an extended period of time. Whilst, this is appropriate for a short period of time the assumption may not hold for large-scale metabolomics data over months or years and currently there are no existing methods to account for normalization over an extended period.

**Table 1.**
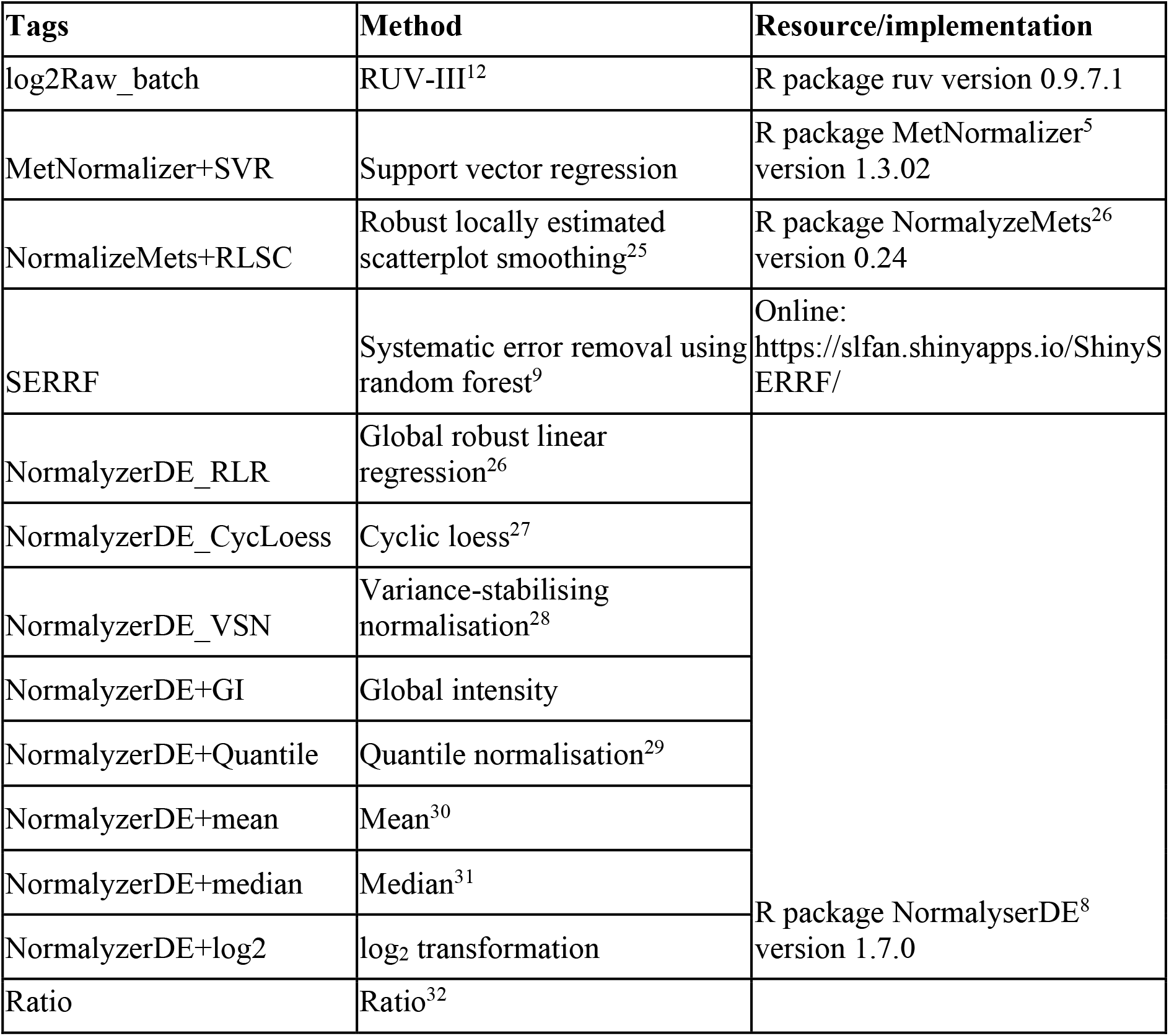
List of existing normalisation methods.

In this paper, we present a novel experimental sample arrangement strategy to embed biological sample replicates throughout large scale experiments to facilitate the estimation of unwanted variation within and between batches with RUV-III^12^, which we will refer to as RUV in this paper. We propose a novel hierarchical approach to removing unwanted variation by harnessing information from sample replicates embedded in the sequence of experimental runs/batches and applying signal drift correction with robust linear or non-linear smoothers. An in-house targeted metabolomics study was performed on a hospital-based cohort of patients with atherosclerosis (BioHEART- CT) was conducted based on the proposed sample arrangement strategy, and we utilise this to assess the normalization on a number of criteria including retention of biological signal, low variability among replication, and reproducibility of results in comparison to other existing methods. The hRUV method is accessible as an R package and also as a shiny application at https://shiny.maths.usyd.edu.au/hRUV/.

## Results

### Replicate arrangement strategy in large scale metabolomics study

We developed a series of technical replications designed as a framework to enable effective data harmonization in large cohorts studies or studies over extended periods of time. Our design includes three types of replicates within each 8×12 = 96-well plate, here called batch. These are the (i) classical pooled QC samples, (ii) single sample replicates in each row of a plate from a random selection of non-replicated samples in previous rows which we call ‘short replicates’, and (iii) five randomly selected non-short replicated samples from each plate are replicated to the next plate, which we call ‘batch replicates’. Fig. 1a and b illustrates a schematic layout of the sample replicates design.

**Fig. 1.**
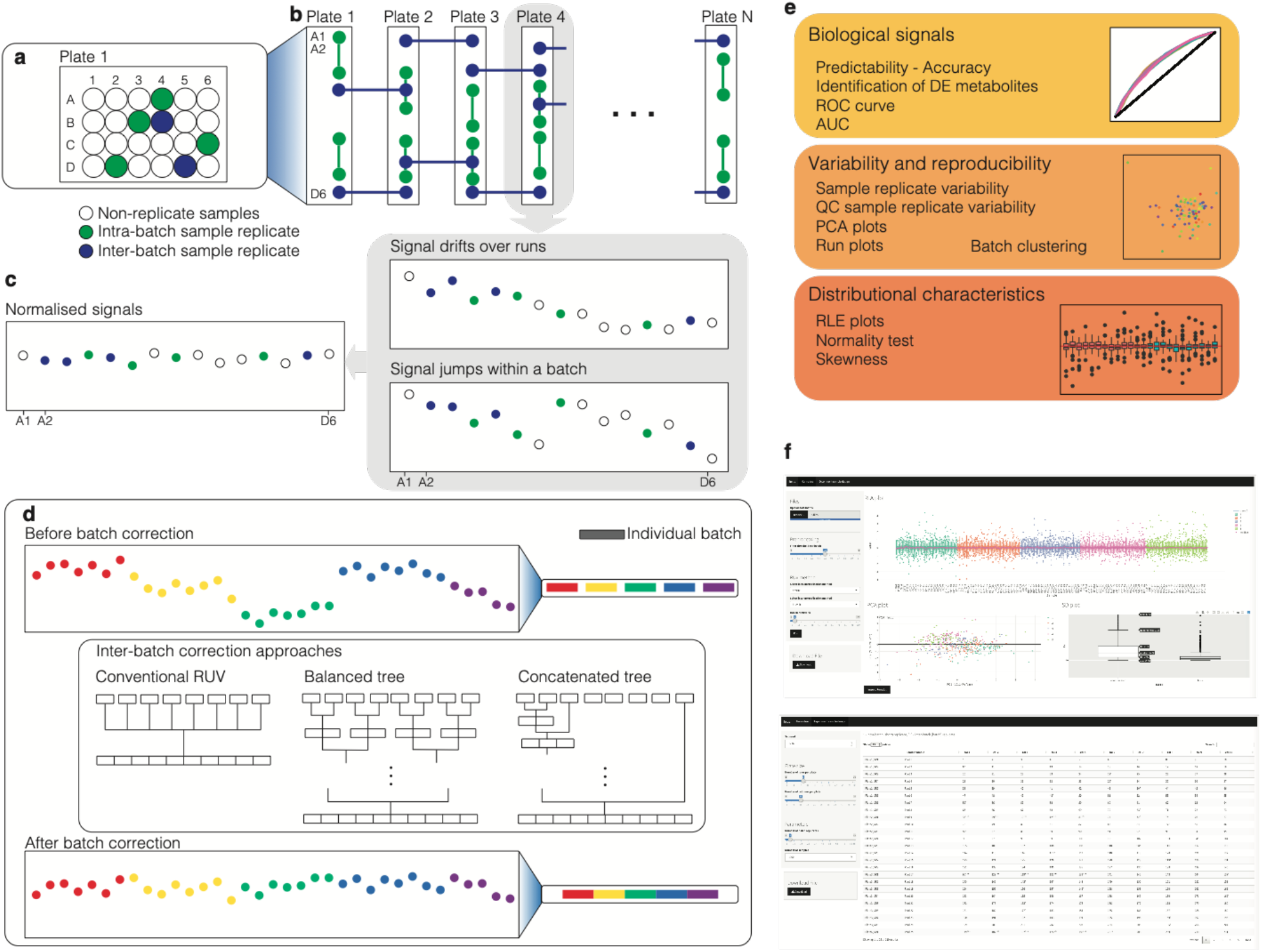
Schematic overview of the hRUV pipeline. **a** A schematic illustration of the plasma sample arrangement for the experimental plate where intra-batch sample replicates (green circles) and inter-batch sample replicates (blue circles) are embedded. **b** A schematic illustration of overall sample replicate design and arrangements. **c** Continuing the color scheme from **b**, two illustrative run plots requiring intra-batch correction, with signal drift and other variations illustrated in the grey boxes. **d** A demonstration of signal variation before and after inter-batch correction in hRUV. A common approach to inter-batch correction is illustrated as conventional RUV and the proposed hierarchical approaches are illustrated. **e** A list of evaluation criteria to assess hRUV performances grouped into categories of biological signals, variability and reproducibility and distributional characteristics. **f** A screenshot of the user-friendly shiny application.

The classical pooled QC consists of a mixture of 10 μL of each of the 1,002 samples, pooled together into a single tube. The pooled QC sample was aliquoted and frozen, and a fresh aliquot was thawed for each batch to minimize the impact of repeated freeze thaw cycles. The spacing of the technical short replicates approximately 10 samples apart capture variation within a short time (approximately 5 hours, based on 30 minutes of run time per sample). This is a good measure of the variation of the metabolomics experiment. In contrast to pooled QC samples, where one sample is repeated many times, short technical replicates are duplicates of different samples; this increase in heterogeneity of samples for the estimation of unwanted variation is more robust compared to estimation with pooled QC. Finally, the batch replicates measure the variation that occurs across different batches. These replicates are typically 60-70 samples apart, capturing variation over a longer time period of 48-72 hours.

This design was used to generate data for a large metabolomics study consisting of 1,002 individuals from the BioHEART-CT biobank and quantification of 100 metabolites per individual. The exact sample designs are given in Supplementary Table 1. In total we had 164 replicates from one pooled QC sample, 230 duplicates from samples across 15 batches, and 140 ‘batch’ duplicates from 70 samples. As expected, variation between replicates within a batch tends to be smaller compared to replicates between batches, as demonstrated in Supplementary Fig 1. A shiny application was developed to enable easy generation of the replicate design upon input of the desired plate size and the desired number of inter-plate replicates. The experimental design with appropriate numbers and assignment of replicates inserted is then exported as a Comma Separated Values (CSV) file. The extra replicate tubes were prepared during the aliquoting and inserted into the appropriate positions during sample processing.

In this current study, a 96 well plate was the natural unit to define as a batch, but in practice, one could select any number of runs (e.g., 100, subjects to the tray capacity of the autosampler) as the unit for a batch. The notions of short and long (batch) replicates can be applied to any batch size to assess variation over a variety of distances.

### A novel hierarchical method to remove unwanted variation (hRUV) in large scale omics experiments

To enable effective data harmonization across large cohort studies or across an extended period of time, we propose a novel hierarchical approach to correct for the unwanted variation between smaller subsets of batches individually, and to sequentially expand to the next set of batches. The two key components of the hRUV can be summarized as follows: (i) signal drift correction within batches with a robust smoother that captures the irregular patterns affecting each metabolite; and (ii) a scalable hierarchical approach to removal of unwanted variation between batches with the use of carefully assigned sample replicates.

The signal drift within each batch was corrected using a robust smoother that captures the trends visible by run order. We explored linear (robust linear model) and non-linear (local regression) model fitting smoothers to capture and remove the run order effects in the data. This is because, due to their chemical and physical properties (Supplementary Fig 2a), each metabolite is affected differently across runs within each batch. These unique changes in signal for each metabolite need to be treated separately.

The concept of sequential batch correction is introduced here to enable scalability for large scale cohort studies. This is a clear contrast to the conventional data integration for normalization that involves estimating unwanted variation across all batches as whole (Fig. 1d). Supplementary Fig 2b shows the inter-batch variation and the differences in the corresponding adjustment factors over time, highlighting the need for dynamic normalization. To this end, we propose two tree structured approaches to estimating the different forms of unwanted variation across a large-scale cohort study, and to dynamically modify the batch effect removal across time. Fig. 1d illustrates the two approaches: balanced tree and the concatenating tree. The balanced tree approach requires log_2_(*n*) RUV adjustments to deal with *n* batches, while the concatenating tree approach requires *n*-1 RUV adjustments. The concatenating approach requires more computation than the balanced tree approach but has an advantage when future integration with new batches is necessary. For once the initial batches are normalised, the additional RUV adjustments are needed are only as many as the number of new batches. While the balanced tree approach is quicker for large *n*, if *m* new batches are introduced in the future, the data will require additional log_2_(*n+m*) RUV adjustments from individual batch level.

The details of hRUV are included in the Methods section. The final output of hRUV is a single normalised and batch-corrected matrix with all input matrices merged and ready for downstream analysis.

### Implementation of a smoother and RUV with sample replicates enables effective adjustment of within plate variation

We assess the performance of signal drift correction by comparing the results of smoothers against themselves and against the commonly used approaches (see Methods). Here, we applied both linear and non-linear smoothers to two sets of sample types; pooled QC samples, or all biological samples within a batch. In general, all four adjustment approaches (loess, rlm, loessSample, rlmSample) give adjusted values that have effectively removed any signal drift associated with experimental run order (Supplementary Fig. 3d-f). The intra-batch correction with all biological samples performs comparably to adjustments performed with pooled QC samples (Supplementary Fig. 3). This suggests a possible reduction in pooled QC samples during experimental design, and thus reducing the total run cost.

In addition to using robust smoothers, the use of RUV with short replicates within each batch after a robust smoother further reduced the sample variations as demonstrated in Supplementary Fig. 2c. Thus using a robust smoother and RUV with short replicates provides effective removal of various unwanted intra-batch variations (Fig. 2) and highlights the value of intra-batch sample replicates.

**Fig. 2.**
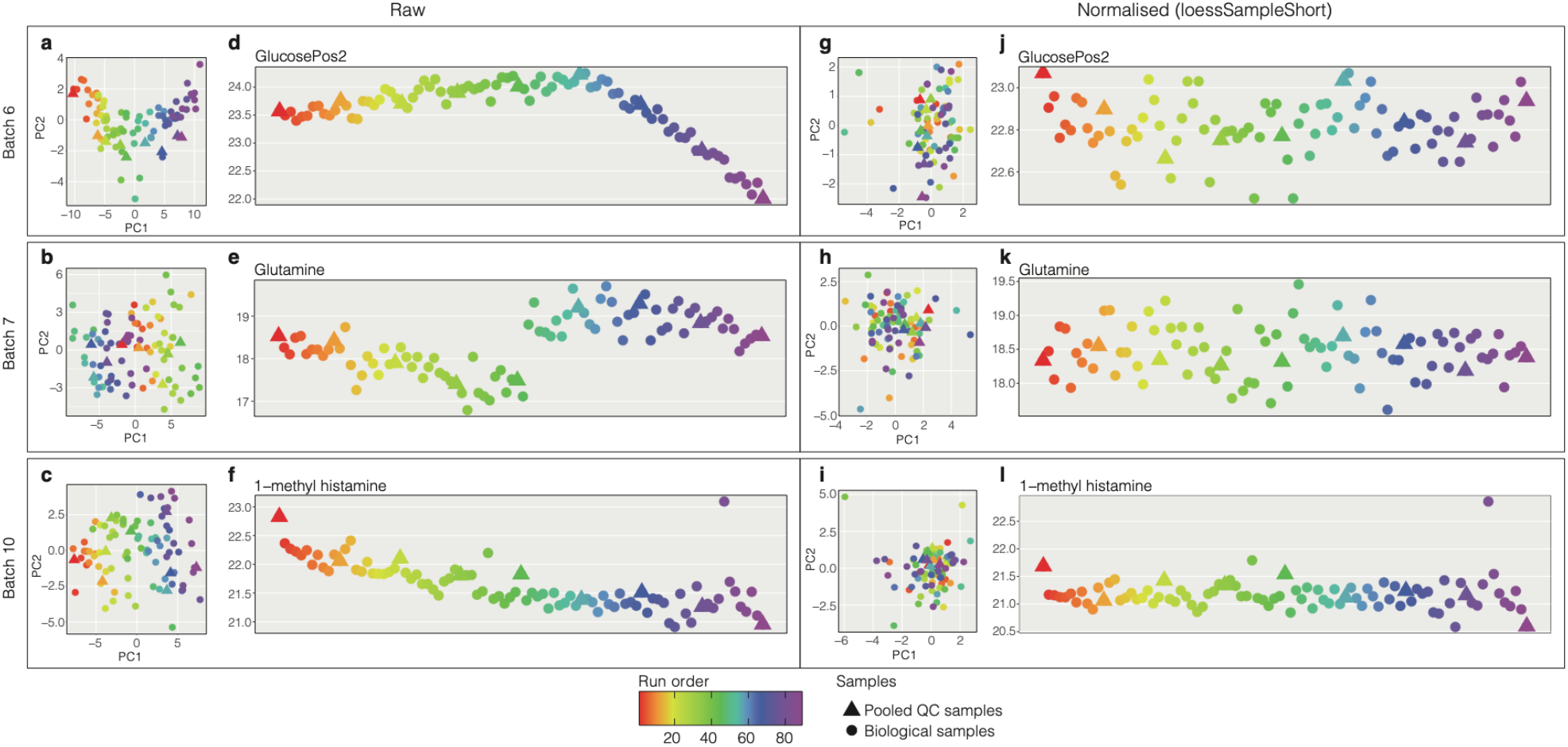
Examples of typical technical variations and signal drifts within each batches for different metabolites and comparison to normalized data. (Note vertical scale change.) **a-c** PCA plots of raw data in batches 6, 7 and 10 respectively, each marker is coloured by the sample run order. **d-f** Run plots exhibiting metabolite signal drift for glucosePos2, glutamine, and 1-methyl histamine respectively, for their respective batches. **g-i** PCA plots from the same batches as **a-c** respectively with intra-batch normalised data using loessSampleShort method. **j-l** Run plot for the metabolites and batches illustrated in **d-f** but with intra-batch data normalized using loessSampleShort method.

### hRUV is more effective in removing unwanted variation compared to other existing methods

Across an extended period of time, there are many different types of unwanted variation. Figure 3a shows that across 1,000 samples, we observe constant or irregular signal drift or abrupt jumps in signals. The run plots (Fig. 3a) illustrate the removal of technical variations introduced between batches and from run time effects for the metabolite glutamate.

**Fig. 3.**
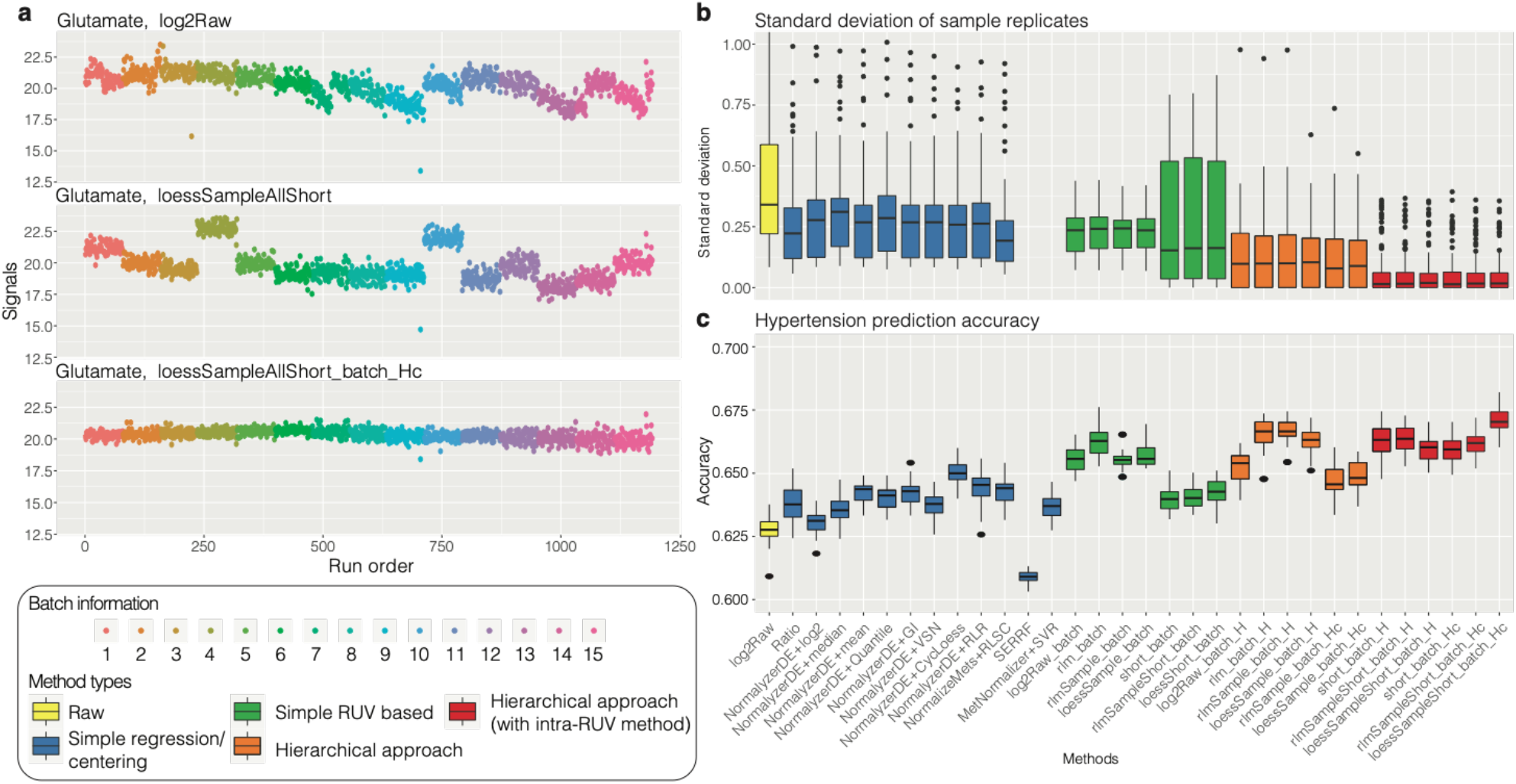
Key assessments of hRUV performances. **a** A run plot of raw, intra-batch corrected and final hRUV normalised data in all 15 batches of the BioHEART-CT cohort. The x-axis indicates the run order, the y-axis indicates the signal of glutamate, and samples are coloured by the batch numbers. **b** Boxplots of all sample replicate standard deviations, where lower values indicate better performance. The boxes are colored by the approach taken to normalize the data. The y-axis of the plot is restricted to a range between 0 to 1 to highlight the differences between the majority of the methods. SERRF and MetNormalizer+SVR’s median sample replicate SD was greater than 1 and thus is not shown. **c** Boxplots of hypertension prediction accuracies for all methods. Higher prediction accuracy indicates better performance. The coloring of the boxes are consistent with that in **b**.

Next, in comparison, we note that the hierarchical RUV normalisation was better at removing unwanted between-batch variation than the original single-step RUV. We compared the standard deviation (SD) between all sets of replicates, with lower values indicating better performance as the replicates should theoretically be identical. Figure 3b highlights lower SD between hierarchical normalization methods (colored in orange and red) and single-step ones (colored in blue and green), suggesting that the hierarchical approach is more effective in batch-correction across extended periods of time. Additionally, hierarchical approaches following intra-batch RUV (colored red) showed even lower sample replicate variation.

### hRUV retains biological signals and outperforms existing normalization methods

To examine the extent to which our method removes only unwanted variation and retains known biological signals, we performed supervised machine learning to illustrate our ability to identify known biological signals for disease prediction. Here we have chosen hypertension as a response variable and performed supervised machine learning classification to detect hypertension status from metabolomics abundance. We anticipate that a normalization method that retains biological signals has a higher classification accuracy. The differential expression (DE) analysis to identify corresponding biomarkers (DE metabolites) measures the interpretability of the signal.

We observed that the average accuracy of hierarchical based normalization methods were generally higher compared to one-step methods (Fig 3c). The ‘loessAllShort_batch_Hc’ method showed the best performance in prediction accuracy. This approach first adjusted for signal drift by fitting a loess line through all the samples and RUV corrected with short technical replicates within a batch, followed by applying RUV in a hierarchical fashion using batch technical replicates.

Proline, valine, cyclic adenosine monophosphate and dimethyl guanidino valeric acid are metabolites known to be associated with individual’s hypertension^13–15^. We observed all these metabolites to be significantly differentially expressed in association with the hypertension status of over 1,000 patients only in hRUV normalized data. Other methods were only capable of identifying subsets of these metabolites.

In general, we found that hRUV performed favorably in terms of maintaining strong biological signal and reducing unwanted variation such as signal drift and batch specific noise in this large study (Fig. 4). Our evaluation metrics capture the trade-off between these two broad objectives. hRUV manages the trade-off between removing unwanted variation and retaining known clinical features of interest. We observed that the ratio, SVR, SERRF and RLSC methods have removed batch effects and reduced sample and pooled QC replicate variance, but as a trade-off these methods result in a loss of biological signals, as evident by the low AUC and prediction accuracy values. Visualizing all these quantities on a heatmap, we find that hRUV methods are ranked in the top 5-10 in most of the evaluation criteria (Fig. 4). The hRUV normalized data show the least variation across the different types of replicate samples and correctly removed batch driven technical noise, whilst maintaining a strong biological signal (Fig. 3b and Supplementary Fig 5a-b).

**Fig. 4.**
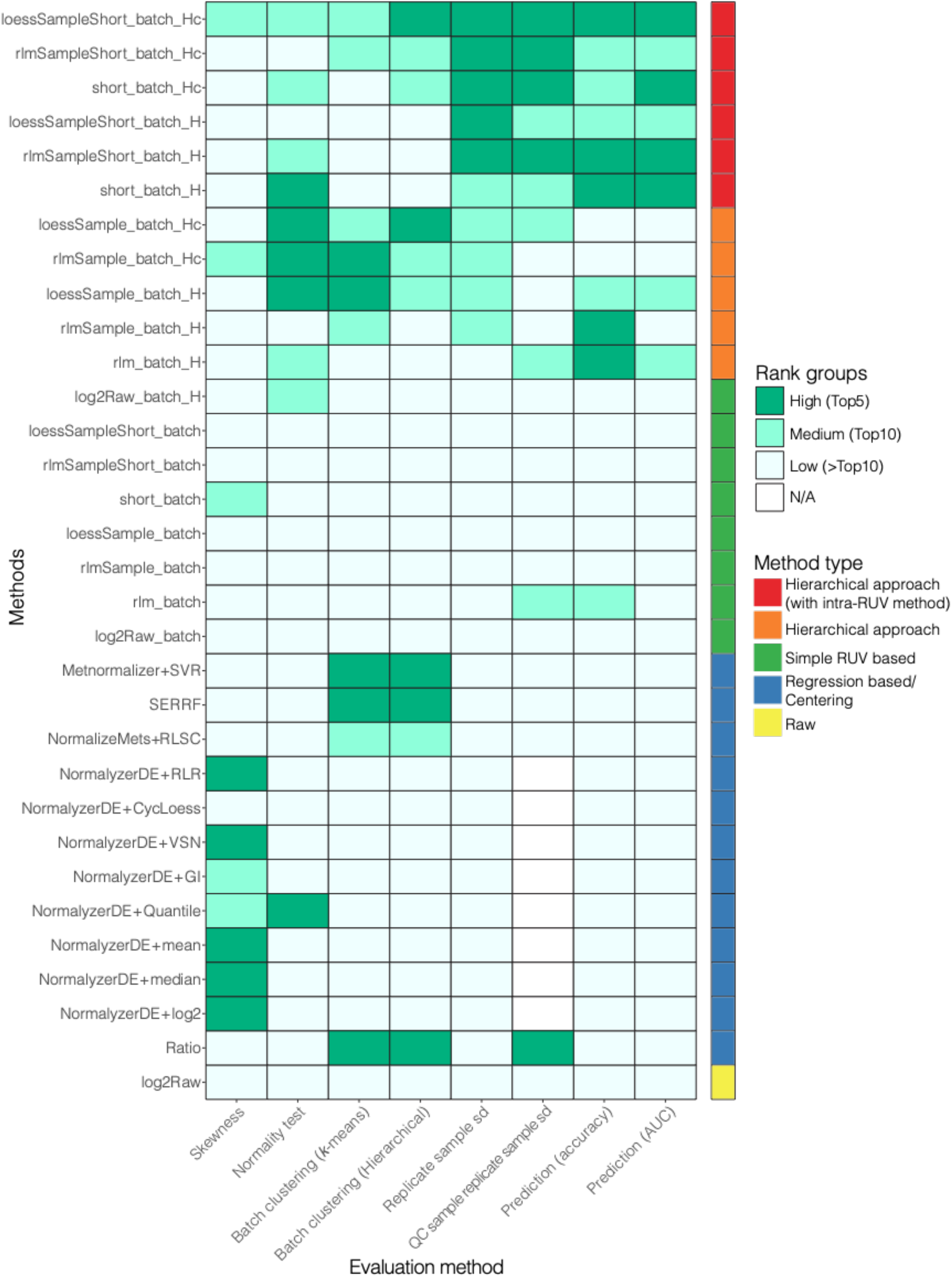
Heatmap of rankings in all evaluations criteria. The y-axis represents all the methods explored in this study and the x-axis represents all the qualitative evaluation metrics used for evaluation of the integrated data. In each category, the evaluation scores are ranked and categorised into 3 groups, high, medium and low. The colored bar on the right indicates the categorized method approaches consistent with **Fig. 3b-c**.

### hRUV is robust to the key decisions and the types of hierarchical structure and choice of negative controls

We have investigated a number of parameters under the hRUV framework, including the various kinds of technical replication, types of negative control metabolite and types of hierarchical structure. We have explored different combinations of replication including pooled QC, inter-batch, and intra-batch replicates. We found that corrections with only pooled QC sample replicates over-estimates the unwanted variation and thus removes biological signals from the data, see Supplementary Fig 6. This highlights the value of using sample replicates as opposed to the pooled QC samples in the estimation of intra- and inter-batch unwanted variation.

In contrast, the two hierarchical approaches in our hRUV show only a small differences in many of the evaluation measures. Both the balanced trees and the concatenating approaches performed adjustments between two sets of batches with 5 inter-batch replicates at each layer. As summarized in Fig 4, the normalization results are very similar between the two types of tree structure.

We explored several approaches to obtaining negative controls for RUV, including a data driven iterative procedure to select stable metabolites and the selection of all metabolites, and saw little difference between the normalised data outputs. We saw that the use of sample replicates demonstrated the greatest impact on the final normalized data (Supplementary Fig. 7).

## Discussion

In this manuscript, we introduce a design strategy and present hRUV, a novel hierarchical approach to remove unwanted variations and batch correct for large scale metabolomics data, where there is a substantial unmet need. We illustrated the performance of this approach using metabolomics data derived from over 1,000 patients in the BioHEART-CT biobank that was run over 15 plates across 44 days.

The careful arrangement of sample replicates on each plate is an important design consideration for large-scale mass-spectrometry studies. Here we believe that systematic arrangements perform better with hRUV normalization than fully random ones. In our current design, the samples to be replicated were selected randomly from the previous plate and the corresponding repeats (batch replicates) were placed at the start of the current plate. Ideally we’d expect to select samples with this strategy whose positions were evenly distributed across the plate, but it is possible by chance to select samples whose positions are from only the first half or only the second half of the previous plate (Supplementary Table 1). This unintended clustering of selected samples was observed between batches 6 and 7, and also between batches 13 and 14, where replicates are selected only from the second half of the previous plate. This limits our ability to capture the unwanted variability across the whole plate, and as a result, we saw a slight shift in signal between these two batches for selected metabolites (Supplementary Fig 5c).

While the proposed hRUV algorithm expects data without missing values, this is often not possible in large-scale metabolomics data due to the nature of the mass spectrometry technology pragmatic issues related to real world clinical studies. To this end, we include an option for users in which the missing values are first imputed prior to applying hRUV and the missing values can be replaced back after hRUV integration. This allows many more sparse metabolites to be incorporated for downstream analyses which is an important aspect in large-scale metabolomics studies and may improve our chance of identifying novel metabolites from the data.

The negative controls are used in RUV to estimate the unwanted variation. The challenge with metabolomics is that the signals of the metabolites are dependent on their individual chemical properties^4,11^ and thus the selection of appropriate negative controls to correct for batch effect is a challenge. Whilst hRUV function accepts a user-defined set of negative controls, in our exploration of data driven negative control metabolites compared to all metabolites as negative control, we have found no significant differences between the two approaches in removal of unwanted variation and utility of inter-batch sample replicates were more effective for batch correction (Supplementary Fig. 7).

In summary, hRUV uses sample replicates to integrate data from many batches in large-scale metabolomics studies. We show the value of suitably located sample replicates for estimating unwanted variation and guiding the design of future studies. While several other existing methods exist to correct large numbers of batches for intra-batch signal drift and inter-batch unwanted variation, hRUV performs consistently better than them in retaining biological variation whilst at the same time removing unwanted variation within and across batches.

## Methods

### Clinical samples

The samples used were from the BioHEART-CT discovery cohort, which has been described in detail previously^**16**^. The study was approved by the Northern Sydney Local Health District Human Research Ethics Committee (HREC/17/HAWKE/343) and all participants provided informed, written consent. Briefly, patients undergoing clinically indicated CT coronary angiogram for suspected coronary artery disease were recruited from multiple sites in Sydney, Australia. Blood samples were taken at time of recruitment, and after appropriate processing, plasma samples including replicates were aliquoted and stored at −80°C until analysis.

Metabolites were prepared as previously described^17,18^. In brief, 10μl plasma was mixed with 90μl HILIC sample buffer, an acetonitrile: methanol: formic acid mix (75:25:0.2, v:v:v). The resulting mixture was vortexed and spun at 14,000 rpm for 20 minutes to remove plasma protein. The metabolite containing supernatant was then transferred to a glass HPLC sample vial and resolved on an Agilent 1260 Infinity HPLC System, and m/z was determined by Qtrap5500 (Sciex)^17,18^. Each sample was eluted over a 25 minute period, and each batch of samples took 40 hours to complete. A total of 15 batches were completed over 44 days.

### Technical replicate design

For each 96 (8 by 12) well plate, in order of run by rows, the first three wells were populated with three pooled QC samples, then a single pooled QC sample was repeated after every 10 runs. For each row of the plate from the second onwards, we randomly selected a single biological sample from the previous row to be replicated at random position in a row (short replicate). For each plate, after the first three pooled QC samples, a random selection of 5 biological samples from the previous plate was repeated (batch replicate) and short replicates are embedded at each row. All randomization was performed using the sample function in R. The replicate design is available as a function `plateDesign` in the *hRUV* package, and also in our shiny application http://shiny.maths.usyd.edu.au/hRUV.

### Pre-processing of metabolomics data

Metabolite elution characteristics were pre-determined using pure standards. Metabolite abundance peaks were integrated using the area under the curve for calibrated peaks from MultiQuant (SCIEX), with manual adjustments to the curves when necessary. This ensures the consistency of all the peaks integrated. The m/z signals were log_2_ transformed and metabolites that were not present in at least 50% of the samples were filtered out. Missing values were imputed using k-nearest neighbour with default parameters implemented in *DMwR2*^**19**^ package in R. We examined the three consecutive QC samples embedded at the start of each plate and removed any outlying measurements.

### Hierarchical approach to removal of unwanted variation (hRUV)

The hRUV algorithm was designed for experiments with a large number of batches, and consists of two key components, including (i) within batch signal drift adjustment with robust smoothers and RUV; and (ii) the adjustment of the datasets with unwanted variation using an RUV in a hierarchical approach. The main inputs to hRUV consist of a list of raw signal matrix, with rows corresponding to metabolites and columns to samples as `SummarizedExperiment`^20^ objects in R, a specific intra- and inter-batch normalisation method, structure of the tree and the parameters for RUV. hRUV performs repeated RUV procedures to sequentially adjust the data over a large collection of batches, with the number of unwanted variation factors (*k*) defaulting to 5.

#### Part I: Signal drift adjustment within a batch

In the present setting, batch refers to one 96-well plate. However, this can be any pre-specified number of samples.

##### (i) Standard adjustment (ratio)

The signal ratio was calculated by dividing the sample signal to its nearest pooled QC sample run. Let us denote *P*_*L*_ as the early run pooled QC sample at run *L* and *P*_*L+M*_ as the next pooled QC sample in a batch at run *L+M*, where *M* denotes the number of run gaps between *P*_*L*_ and *P*_*L+M*_. Then the signal ratio is defined as follows:

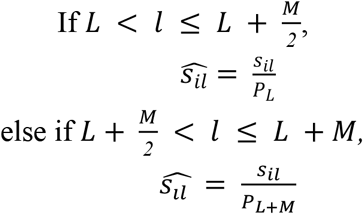

where *s*_*il*_ denotes a signal of a sample with metabolite *i* at run number *l*.

##### (ii) Loess line

A loess line was fitted to all the biological samples for each metabolite within a batch with the default span parameter of 0.75. The differences between the fitted line to the median of each metabolite across all samples per batch were calculated for adjustment of each samples as follows:

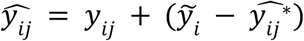

where *y*_*ij*_ represents a *log*_*2*_ transformed signal for sample *j* in metabolite *i* in a batch and 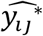 denote a loess fitted value of *y*_*ij*_ and 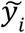 denote the median of *y*_*ij*_ for all *j*. Here, the `loess` line uses the loess function in the *stats*^21^ package.

##### (iii) Linear line

Robust linear model (rlm) was fitted on a *log*2 transformed signal against the run index using the `rlm` function from the *MASS*^22^ package to all the biological samples with a maximum iteration set to 100. The adjustments of each sample were calculated as with to the loess approach where 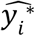 would denote the predicted value of *y*_*i*_.

##### (iv) Loess line fitting with pooled QC samples

Similar to (ii), we fit the loess line to the pooled QC samples only. The adjustment values of each sample were calculated using the predicted values from the model 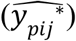.

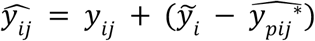

##### (vii) Linear line fitting with pooled QC samples

Likewise, we fit the rlm to the pooled QC samples. The adjustment values of each sample were calculated using the predicted values from the model.

##### (viii) RUV based approaches

We incorporated sample replicates into our design matrix of RUV introduced by Molania et al^12^. These sample replicates are utilised to estimate the unwanted variation as the signals of these replicate samples should theoretically be identical. All metabolites were selected as the negative controls for RUV and the number of unwanted factors to use (*k*) was taken as 5.

#### Part II: Hierarchical batch integration design

##### (i) Balanced tree

The balanced tree approach to normalisation is to perform removal of unwanted variation measured between pairs of different batch groups at different levels of the tree. In this approach, we began by removing unwanted variation between pairs of neighbouring batches. In the next layer of adjustment, we paired the two neighbouring groups of integrated batches (sets of 2 batches) and repeated the process to expand the number of batches per set until the last layer, where we had a single group of all the batches which was normalised, as illustrated in schematics in Figure 1d. For a study with *n* batches, this will requires *log*_*2*_*(n)* RUV adjustments.

##### (ii) Concatenation

As with the balanced tree approach, the concatenating approach performs removal of unwanted variation measured between pairs of batches, but in a sequential progression. We began with the first two batches for batch correction and sequentially introduced the next batch to remove unwanted variation as illustrated in schematics in Figure 1d. For a study with *n* batches, this will require an *n-*1 number of RUV adjustments.

For both balanced tree and concatenation methods, we apply RUV at each layer as follows:

Let us denote by *B* the pair of batches of interest, *M* as the number of metabolites, and S as the number of samples in batches B. The mean adjusted sample *Z*_*mbc*_ can be calculated as:

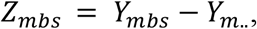

where *Y*_*m..*_is the average expression of metabolite *m* across samples *S* and batches *B* calculated by:

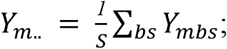

The mean adjusted data *Z*_*S×M*_can be fitted to the model underlying the RUV model, which is formulated as:

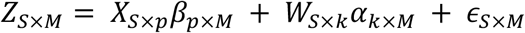

where *X* is the matrix of factor of interest; *p* is the number of factors of interest; *W* is the unobserved design matrix corresponding to the unwanted factors; *k* is the linear dimension of the unwanted factors, which is unknown; ε denotes the random error. Thus the RUV normalised data can be represented as:

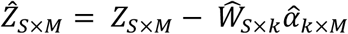

After the RUV, *Y*_*m..*_ is returned back to the mean adjusted RUV normalised data as follows:

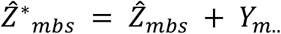

### Data driven negative metabolite selection

We explored adaptive data driven selection of negative control metabolites in comparison to a selection of all metabolites in an RUV method. The adaptive selection was performed by ranking non-differentially expressed metabolites by *p*-values per batch for the hypertension response variable. We utilised differential expression analysis with the *limma*^23^ package (version 3.46) in R.

### Performance evaluation / evaluation metric processing

We evaluate hRUV methods including 13 publicly available metabolomics data normalization methods (Table 1). Details of the method abbreviations is explained in Table 2. These packages were installed either through the official CRAN or Bioconductor website where available, or from GitHub pages. For all 13 existing methods, we used the default settings and parameters as described in the package README or vignette for training each model.

**Table 2.**
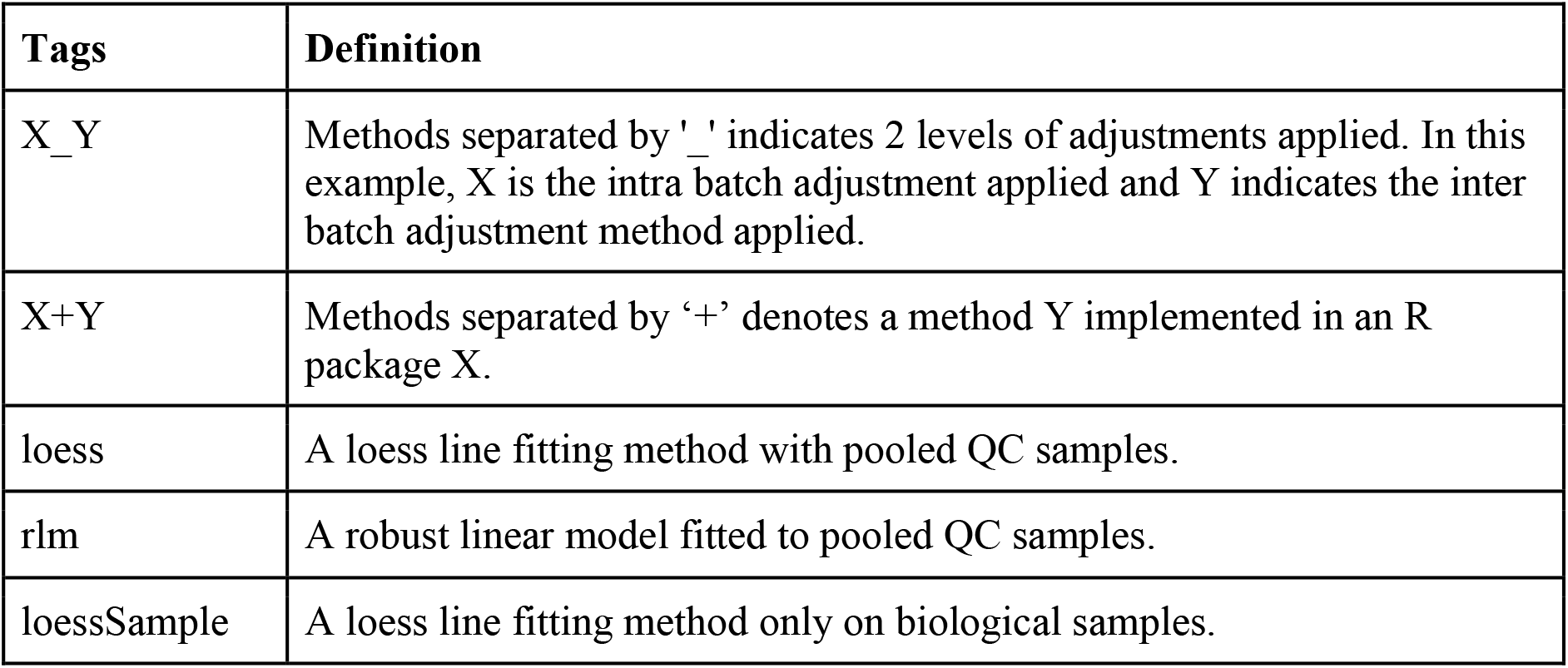
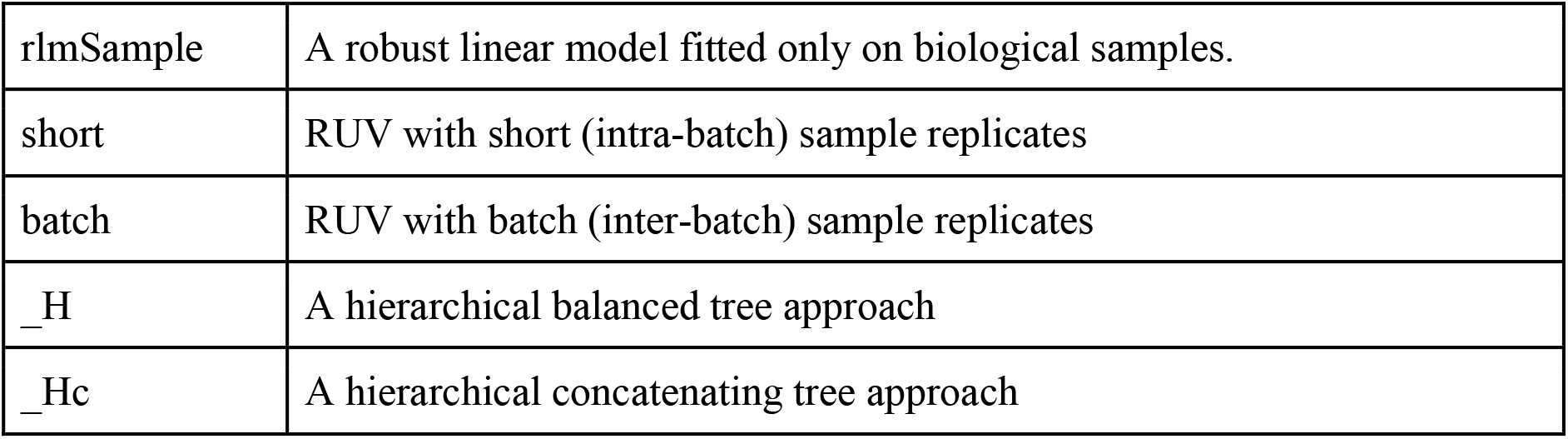
A normalisation method abbreviation dictionary.

### Evaluation metrics and plots

#### (i) Skewness

The skewness of samples were calculated with `skewness` function from *e1071*^24^ package in R.

Let us denote *x*_*j*_ for the non-missing elements of **x**, *n* for the number of samples, μfor the sample mean, *s* for the sample standard deviation, and *m*_*r*_ = ∑_*j*_(*x*_*j*_ − *μ*)^*r*^/*n* for the sample moments of order *r*. The skewness then can be calculated as: Skewness = *m*_*3*_/*s*^*3*^,

#### (ii) Normality metric

The normality tests were performed with Shapiro-Wilk normality test implemented in `shapiro.test` function from the *stats*^21^ package in R.

#### (iii) Predictability with accuracy

To assess the predictability of a normalised dataset, we utilised a binary diagnosis of hypertension as the response variable. This was chosen as it had a reasonably balanced class distribution, as 39% of the cohort had hypertension. We used a Support Vector Machine (SVM) implemented in the *e1071*^24^ package to predict the hypertension status of participants of the study. We measured the average accuracy via a 30-repeated 10 folds cross-validation strategy.

#### (iv) Signal strength with AUC

We use the same prediction model from (iii) and calculate the area under the ROC curve (AUC) values.

#### (v) Standard deviation of replicates (SD replicates)

To demonstrate the variation between the replicate samples after normalisation, for each set of replicated sample, we calculated the standard deviation for each metabolite and visualised the results as a boxplot. A low standard deviation indicates a small variability between the replicates and thus illustrates that the replicates are close to identical.

#### (vi) Clustering by batch (Reduction in batch effect)

To assess the removal of batch effects, we performed unsupervised hierarchical and *k-*means clustering (`hclust` and `kmeans` in *stats*^21^ package in R respectively) where we set the number of clusters *k* to the number of batches. The cluster output is evaluated using adjusted rand index (ARI):

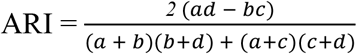

where *a* is the number of pairs of samples partitioned into the same batch group by the clustering method, *b* is the number of pairs of samples partitioned into the same cluster but does not belong to the different batch group, *c* is the number of pairs of samples partitioned into different clusters but belongs to the same batch group and *d* is the number of pairs of samples correctly partitioned into different clusters. A low ARI value indicates lower concordance with the batch information and thus demonstrates removal of batch effect in the data.

#### (vii) Differential expression (DE) analysis of hypertension

To assess the biological signal in the normalised data, we performed DE analysis with the R package *limma*^23^. We identified a set of metabolites with a 5% level of significance and verified their association with hypertension from the literature.

### Diagnostic Plots

To graphically assess whether the normalization method or the choice of parameters of hRUV has effectively corrected the batch effect, we have provided three kinds of diagnostic plots: (1) PCA plots; (2) relative log expression (RLE) plots^19^; (3) metabolite run plots.

#### 1. PCA

PCA plots were generated using all metabolites. We show the first and second principal components.

#### 2. Relative Log Expression (RLE) plot

RLE plots are a useful tool to visualize unwanted variation. RLA plots are boxplots of RLA for each sample, calculated as 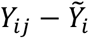, where 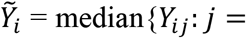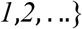, and *Y*_*ij*_ is the log signal value of metabolite *i* in sample *j*. The samples from different batches should have a similar distribution, and the medians of the boxplots should be close to zero if the unwanted variations are removed.

#### 3. Metabolite run plot

Metabolite plots are a useful diagnostic visualisation to visualise the signal drifts. The run plots are a scatter plot of signals for each metabolite against the run order of all the samples. The overall shape of the scatter plot should be a flat horizontal bar. All other shapes of trend in the scatter plot is an indication of a signal drift.

## Supporting information

Suppementary information

## Author contributions

JYHY, GF and JO conceived and designed the study. JY designed the sample replicate framework with input and discussion from TPS, JO, DJ and TK implemented the design. The BioHEART study was designed by GF and the metabolomics assays were performed by OT and JP supervised under YCK and JO. OT and JP processed and curated the data with input from JO, YK, JY. GF, SV & KK provided the clinical and pathology input and guidance with BioHEART clinical data. JY, PY and TK led the hRUV method development with input from TPS. JY and TO lead the evaluation and data analysis with input from GF, JO, TP, PY, and OT. TK implemented the R package with help from PY and JY. All authors wrote and reviewed the manuscript have approved the final version of the manuscript.

## Acknowledgements

The authors thank all their colleagues, particularly at The University of Sydney, School of Mathematics and Statistics, Charles Perkins Center, Kolling Institute and Royal North Shore Hospital for their support and intellectual engagement. The following sources of funding for each author, and for the manuscript preparation, are gratefully acknowledged: Australian Research Council Discovery Project grant (DP170100654) to JYHY; Judith and David Coffey Lifelab Scholarship to TK. KK is supported by an Australian Commonwealth Government Research Training Program Stipend Scholarship; SV is supported by a University of Sydney Postgraduate Research Scholarship funded by Heart Research Australia. GF is supported by a National Health and Medical Research Council Practitioner Fellowship (grant number APP11359290), Heart Research Australia, and the New South Wales Office of Health and Medical Research. PY is supported by a National Health and Medical Research Council Investigator Grant (APP1173469).

The funding source had no role in the study design; in the collection, analysis, and interpretation of data, in the writing of the manuscript, and in the decision to submit the manuscript for publication.

## Declaration of competing interests

None

## Data availability

The sample designed is made available on Supplementary Table 1. The mass spectrometry metabolomics data will be available after publication on the Github repository [TBA]. All other data are available from the corresponding author on reasonable request.

## Code availability

The hRUV implementation is available as an R package stored at the GitHub, https://github.com/SydneyBioX/hRUV and as a web shiny application at http://shiny.maths.usyd.edu.au/hRUV.

## Notes

### Competing Interest Statement

The authors have declared no competing interest.

