## Supplementary material for "hRUV: Hierarchical approach to removal of unwanted variation for large-scale metabolomics data": Suppementary information

**Kim et al.**

**Supplementary Figures**

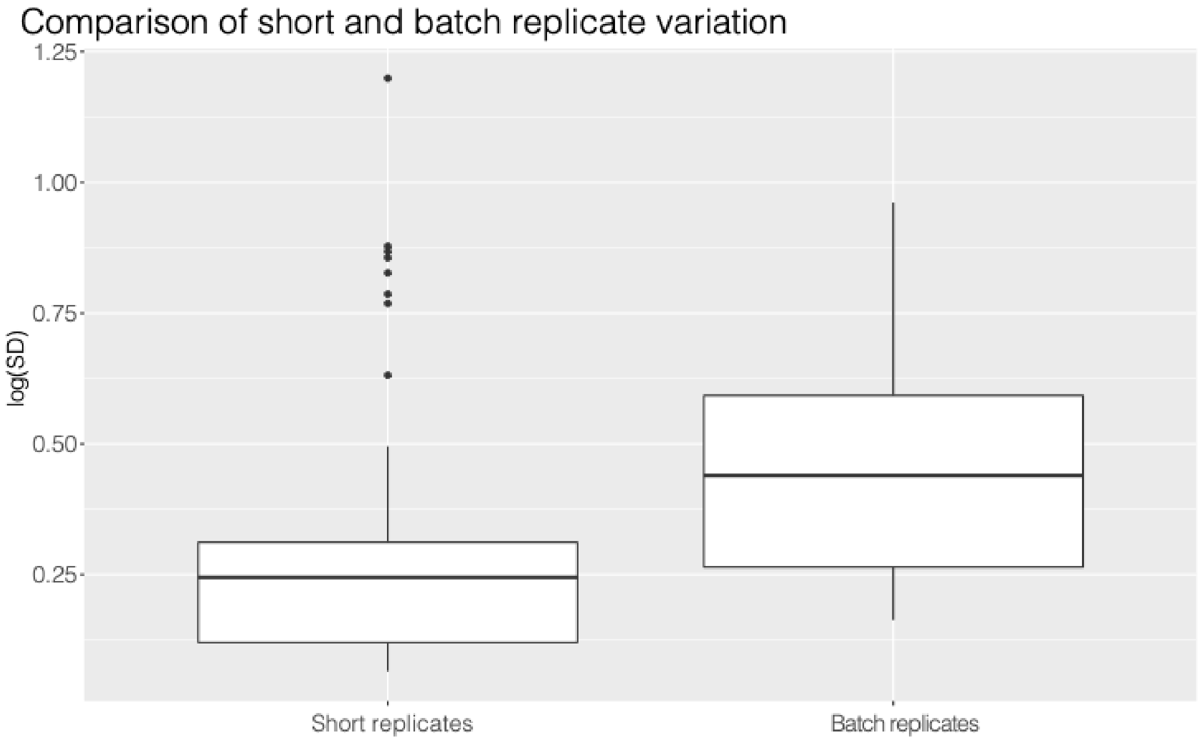


**Supplementary Fig. 1**. Boxplots comparison of intra- (short) and inter- (batch) replicates standard deviations (SD). The y-axis is a natural log transformed SD. The batch replicate samples exhibit higher standard deviations than the short sample replicates.

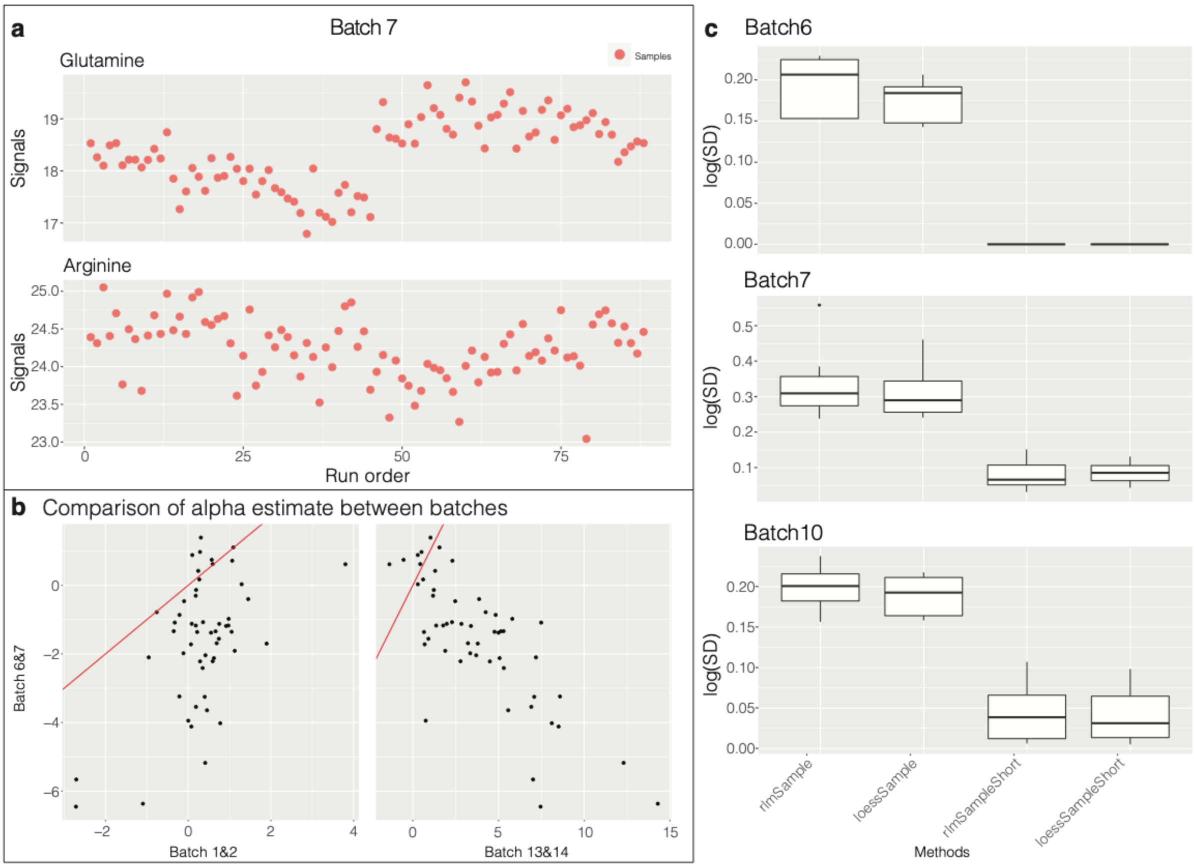


**Supplementary Fig. 2. a** A run plot of raw data values from batch 7 exhibiting signal drift in the glutamine and arginine measurements. **b** A comparison of the first component of the estimated alpha term in the intra-batch RUV correction across two adjacent pairs of batches. The y-axis shows the alpha values estimated from batches 6 and 7. The x-axis shows the alpha values estimated from batches 1 and 2 (left) and also from batches 13 and 14 (right). The red line is y=x. **c** Boxplots of sample replicate standard deviations for the robust smoother with and without RUV using short replicates. The y-axis is a natural log transformed SD.

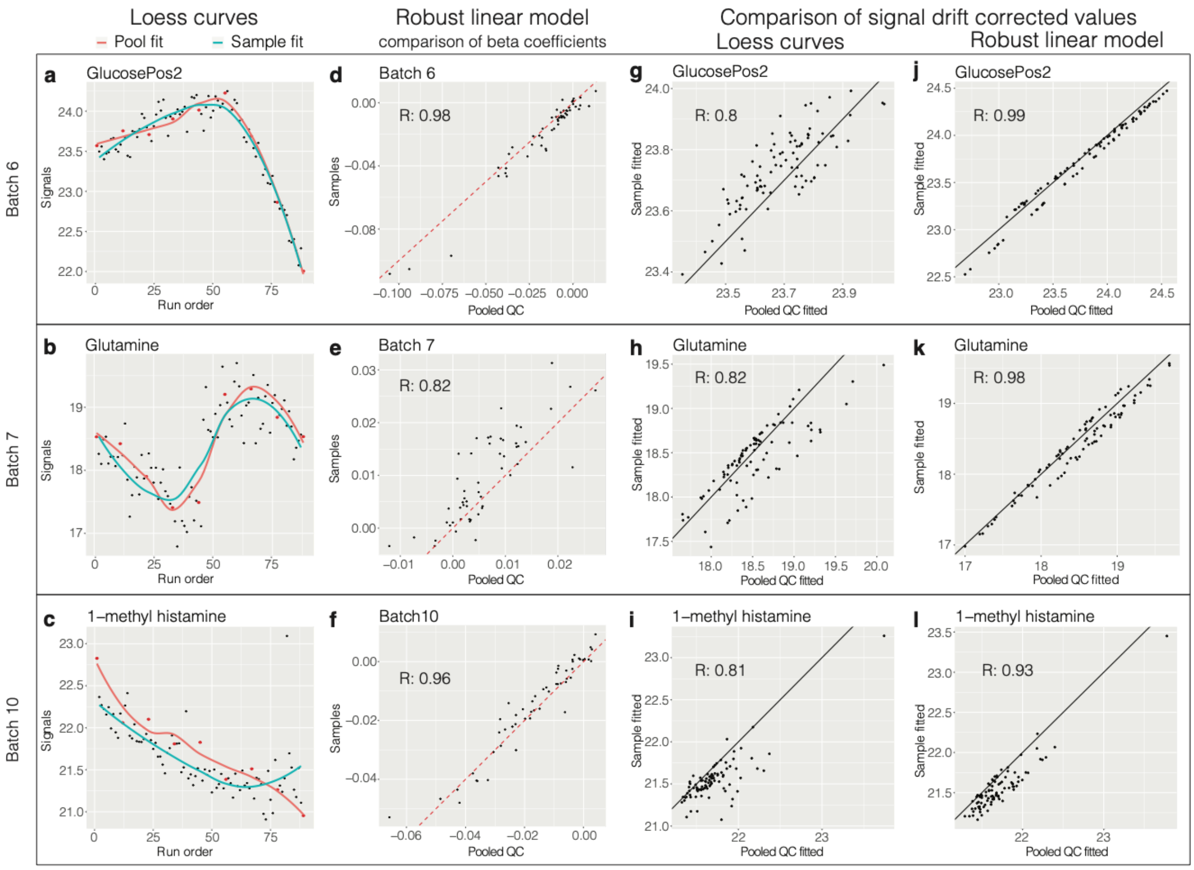


**Supplementary Fig. 3.** Comparisons of pooled QC fitted smoother and sample fitted smoothers. The rows of the panel exhibit data from Batch 6, 7 and 10 samples. The columns exhibit types of comparisons. **a-c** A pooled QC fitted loess curves (red) and sample fitted loess curves (blue) of glucosePos2 in Batch6, glutamine in Batch 7 and 1-methyl histamine in Batch 10 respectively, all against run order. **d-f** Beta coefficients of sample fitted robust linear model (y-axis) against pooled QC fitted robust linear model (x-axis). **g-i** Signal drift adjusted values from sample fitted loess smoother (y-axis) against pooled QC fitted values (x-axis). **j-l** Similarly to **g-i**, with signal drift adjusted values from pooled QC fitted (x-axis) and sample fitted (y-axis) robust linear models.

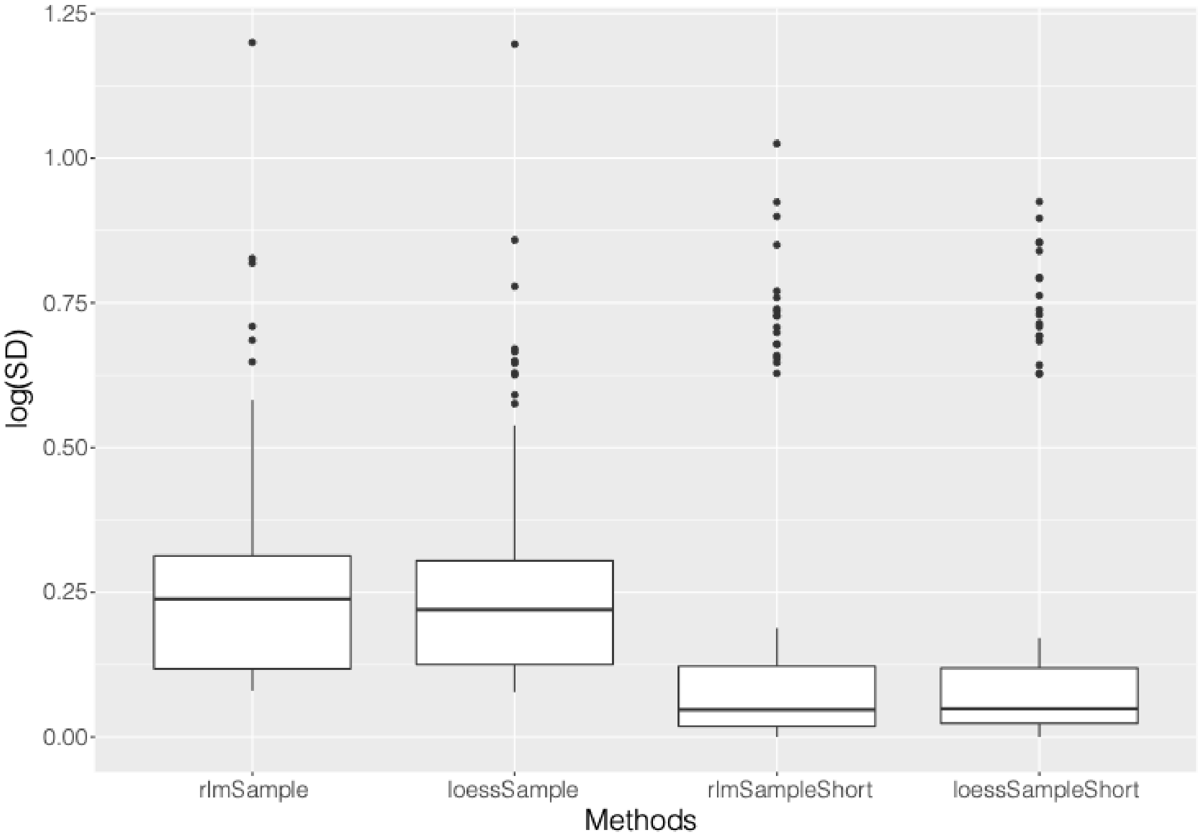


**Supplementary Fig. 4**. A comparison of intra-batch normalisation methods with and without RUV using short replicates. The y-axis is a natural log transformed SD.

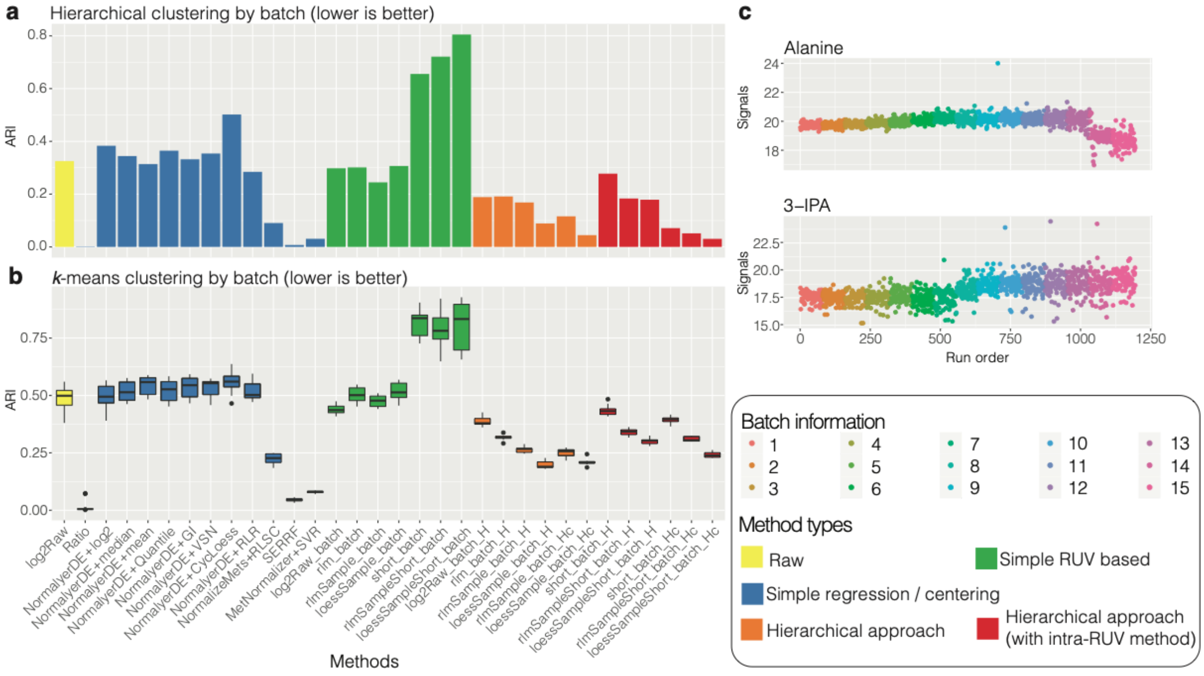


**Supplementary Fig. 5. a** A bar plot of Adjusted Rand Indices (ARI) which compares the concordance of hierarchical clustering to the known batch information. Higher ARIs indicate clusters driven by batch effects; thus lower ARIs indicates better normalisation. **b** Similar to **a**, a boxplot of ARI concordance of 10 permutations of *k*-means clustering to the known batch information. **c** A sample run plot for the metabolites alanine and 3-indolepropionic acid with loessSampleAllShort_batch_Hc normalised data. The samples are colored by batch number.

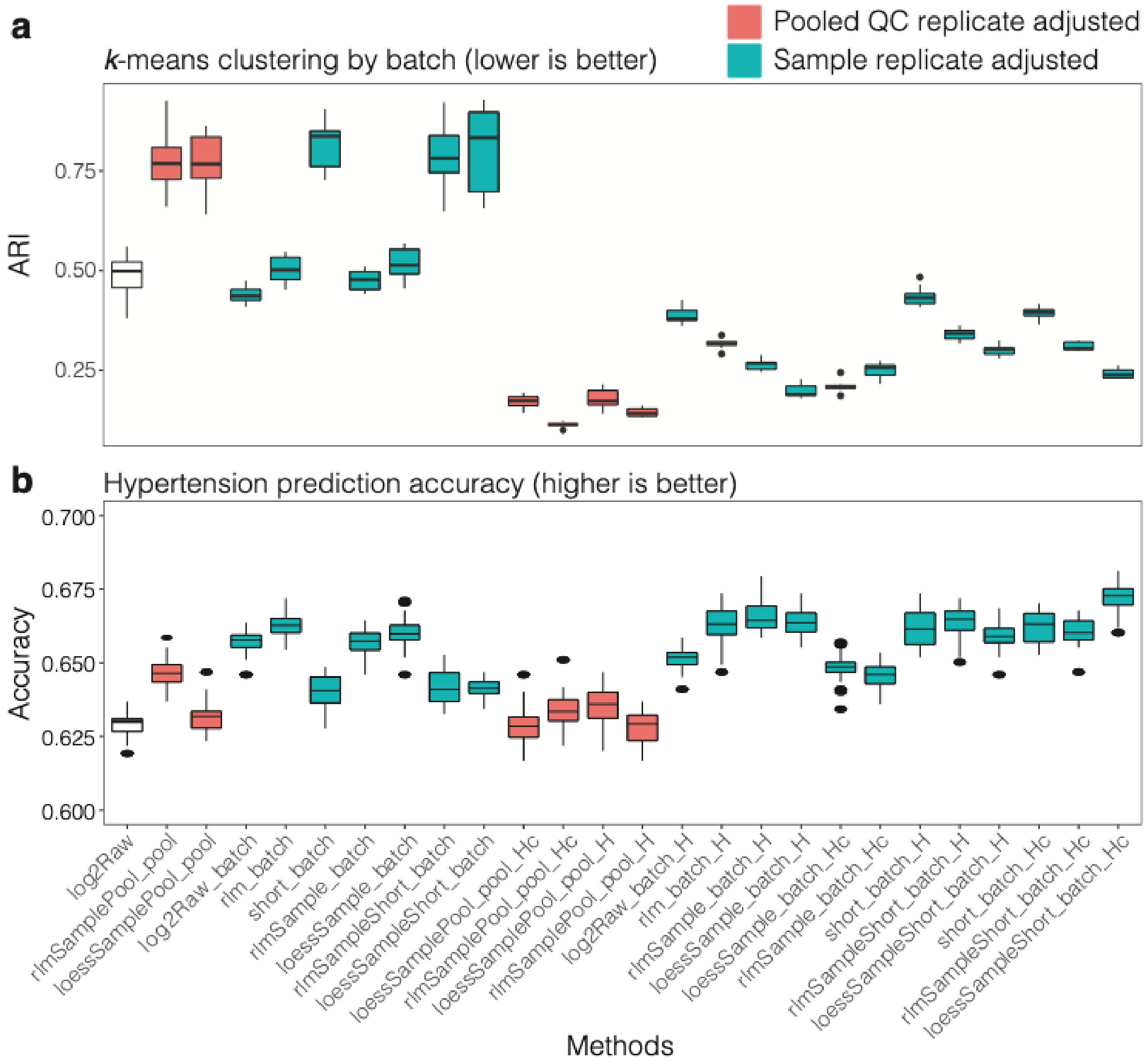


**Supplementary Fig. 6**. **a** Boxplots of clustering ARIs compared to batch with pooled QC replicate adjusted RUV and sample replicate RUV adjusted data. **b** Boxplots of hypertension prediction accuracies for the same methods as in **a**.

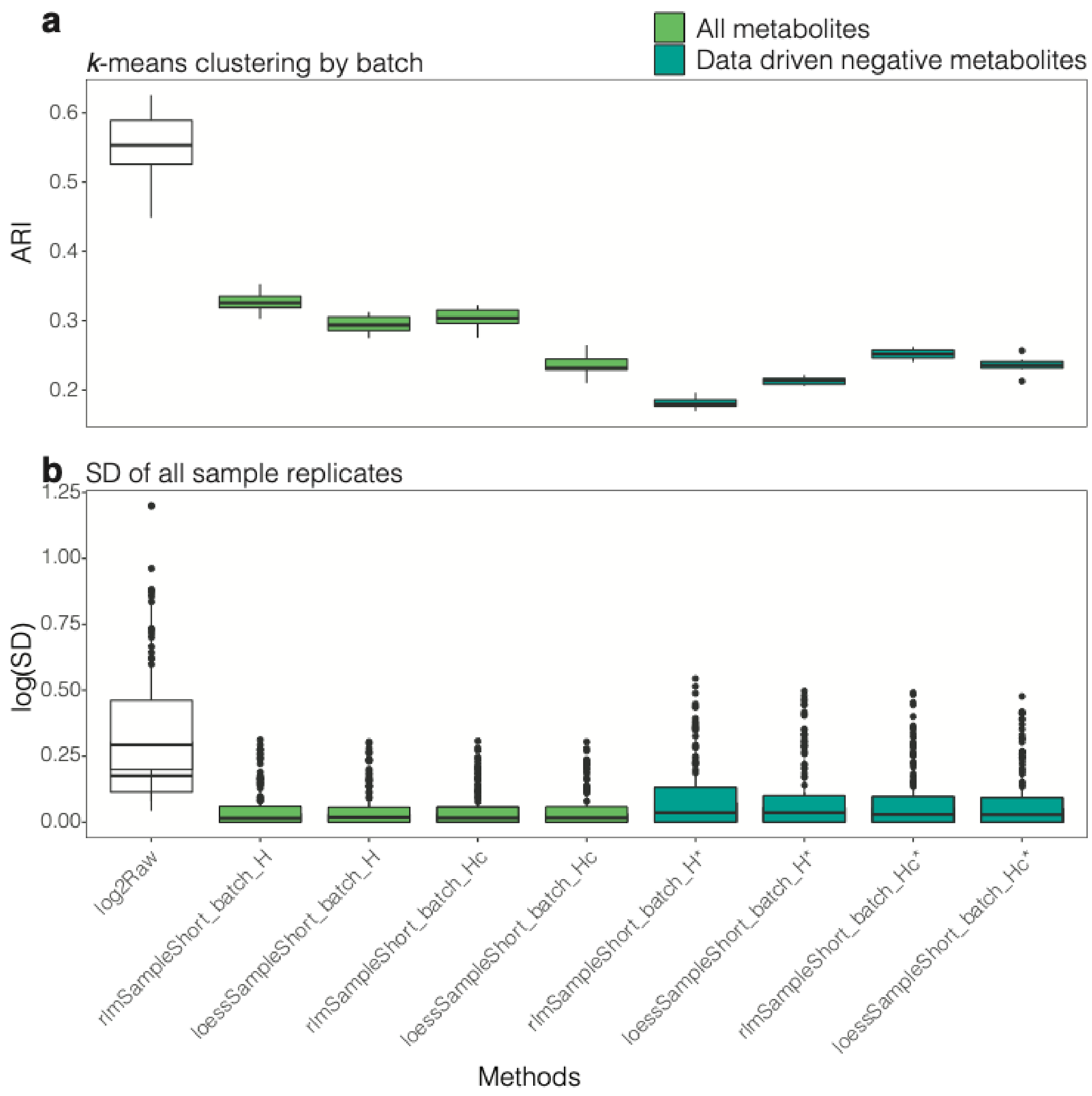


**Supplementary Fig 7**. Comparison of results using different sets of negative controls in RUV. **a** Boxplot comparison of ARIs from *k*-means batch clustering. **b** Boxplots of standard deviations from all sample replicates for the normalisation methods with all metabolites and data driven negative metabolites as negative controls in RUV. The y-axis is a natural log transformed SD.
